# Deconvolution of gene expression noise into physical dynamics of cognate promoters

**DOI:** 10.1101/019927

**Authors:** Ángel Goñi-Moreno, Ilaria Benedetti, Juhyun Kim, Víctor de Lorenzo

## Abstract

When facing recalcitrant pollutants, soil bacteria exploit noise of catabolic promoters for deploying environmentally beneficial phenotypes such as metabolic bet-hedging an/or division of biochemical labor. While the origin of such noise in terms of upstream promoter-regulator interplay is hardly understood, its dynamics has to be somehow encrypted in the patterns of flow-cytometry data delivered by transcriptional reporter fusions. On this background, we have examined the behaviour of the *Pm* promoter of the environmental bacterium *Pseudomonas putida* and its cognate 3-methylbenzoate-responsive regulator XylS under different conditions by following expression of *Pm*-GFP fusions in single cells. Using mathematical modeling and computational simulations we elucidated the kinetic properties of the system and use them as a baseline code to interpret the observed fluorescence output in terms of upstream regulator variability. Transcriptional noise was predicted to depend on the intracellular physical distance between the regulator source (where the e.g. XylS is being produced in the cells) and the target promoter. Experiments with engineered bacteria where this distance is either minimized or enlarged proved the effects of proximity on noise patterns as predicted by the model. This approach not only allowed deconvolution of cytometry data into mechanistic information on the gene expression flow. But it also provided a mechanistic basis for selecting a given level of noise in engineered regulatory nodes e.g. in Synthetic Biology constructs.

## Introduction

The processing of information inside bacterial cells in response to physicochemical stimuli requires the functioning of regulatory cascades to effectively propagate cognate input/output signals. This typically involves multiple steps in which upstream produced transcription factors (TFs) have to interact with downstream promoters, triggering gene expression responses. These interaction events occur stochastically in time rather than deterministically [1–5], leading to very specific and variable noisy signals. The customary view considers this effect as the necessary consequence of random fluctuations of regulatory elements present in short supply in individual cells [6]. However, the range and intensity of expression noise of given promoters appears in some cases as an adaptive trait that frames the dynamic properties of promoter activation [7–9]. The onset of single-cell technologies [10–12] has shed some light on the various mechanisms behind noise generation. A major source of noise in virtually every prokaryotic promoter is the so-called *bursting* effect [13,14], a pulse-like activity that largely results from discontinuous topological changes of DNA caused by the progression of RNA polymerase (RNAP) through transcribed DNA [15,16]. But this default *pulsing* scenario then intersects with the interplay between of the promoter at stake and its specific regulators in response to particular conditions. The outcome of different noise generators *in vivo* is a distribution of fluorescence in single cells that can be followed through cytometry of bacteria bearing e.g. transcriptional GFP fusions [17]. In other words, cell cytometry profiles embody information on the mechanistic origin of the observed gene expression noise. But how to retrace such data to the physical TF-promoter scenario that produces the distribution of fluorescent signals in a population?

The Gram-negative soil bacterium *Pseudomonas putida* mt-2 provides an exceptional model for tackling the questions above. This microorganism is able to thrive in sites polluted with aromatic chemicals [18] e.g. *m*-xylene (*m*-xyl), because of a complex metabolic and regulatory network encoded in its single-copy TOL plasmid pWW0 [19] (Figure 1). One conspicuous feature of this system is that noise of each of the four promoters of the network seems to be exquisitely controlled for deploying a metabolic bet-hedging strategy [20]. This allows a fraction of the cells in a population (but not all) to explore new nutritional landscapes without risking a communal collapse should such *reconnoitre* fail [20,21]. The noise of the *Pu* and *Ps* promoters of the network can easily be explained by the very low number of molecules of their cognate regulatory protein XylR [22]. However, that of *Pm* (which runs the lower operon of the pathway in response to 3-methylbenzoate 3MBz; [23]) is quite puzzling. As shown in Figure 1, this promoter can be activated through two separate mechanisms i.e. either [i] a low intracellular concentration of the cognate regulator XylS bound to its effector, 3MBz or [ii] overproduction of XylS-alone, with no concourse of 3MBz. Logically, when the two circumstances co-occur (i.e. high XylS levels and presence of 3MBz), *Pm* activity can reach very high activity levels [24,25]. Yet, the revealing feature of this regulatory node is that the noise pattern displayed by *Pm* varies dramatically depending on either mechanism, as will be seen in the Results section. On this background we wondered whether the cell cytometry data of transcriptional *Pm*-GFP fusions could be decoded into information on the physical dynamics of promoter activation, including hints on the arrangement of the XylS*/Pm* regulatory node in the cell.

**Figure 1:**
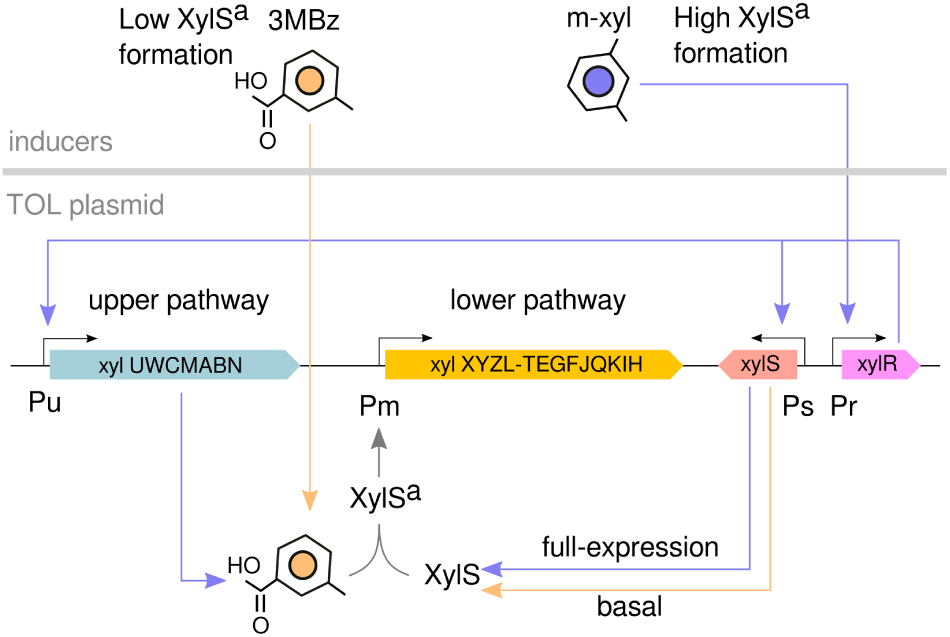
The TOL metabolic and regulatory network borne by plasmid pWW0 of *Pseudomonas putida* mt-2. As shown in the sketch, *m*-xylene is first converted to 3-methylbenzoate (3MBz) through the action of the enzymes encoded by the *upper* TOL pathway, and this intermediate compound is further metabolized into the TCA cycle by the activity of the *lower* pathway. XylR and XylS are transcriptional regulators while *Pu, Pm, Ps* and *Pr* are promoters. The master regulatory gene *xylR* controls expression of both the *upper* pathway and the second transcriptional factor, XylS, which is encoded in a location adjacent to the end of the *lower* operon. In the absence of *m*-xylene, this second regulator XylS is produced at low levels, and it changes from an inactive form to a transcriptionally proficient TF able to induce *lower* pathway expression by activating *Pm.* This regulatory architecture plays a decisive role in the dynamics of *Pm* activation due to the fact that the levels of its cognate activator (XylS), vary depending on the inducer used. In one case, 3MBz activates XylS molecules that are present in low numbers in the cell owing to the leaky expression of the *Ps* promoter. This results in the active form of the protein that we have called XylSa, which is able to bind and activate *Pm.* In the second case, *m*-xylene (*m*-xyl) both causes over-expression of XylS (due to activation of *Ps* by XylR) and intracellular production of metabolic 3MBz (because of the activity of the *upper* pathway operon driven by Pu). Therefore, *m*-xyl leads to a higher concentration of XylSa than externally added 3MBz. This difference is the key feature for decoding *Pm* output, as explained in the text.

The combination of modeling and experimental work presented below shows not only that the noise regimes observed in the *Pm* promoter are the consequence of an specific and steady set of kinetic rates with low XylS-Pm affinity dynamics and high gene expression values. Also, that noise regimes can be changed as needed by changing regulator numbers which, in the non-homogeneous intracellular milieu, ease or not the TFs to reach its target sequence in *Pm*. These predictions were validated in cells engineered to minimize the distance between the source site of XylS and the location of *Pm.* In that sense, the modelling-wet validation pipeline adopted in this work not only allowed deconvolution of flow cytometry data into kinetic details. It also exposed an added biological functionality to the genomic distance between regulatory genes and their target promoters in terms of setting the corresponding output noise, which can thereby be fixed on-demand.

## Results

### Two distinct noise regimes *rule Pm* output

As mentioned above, the activity of the inducible promoter *Pm* of the TOL plasmid (Figure 1) can be triggered by exposing *P.putida* mt-2 to either one of these two inputs: [i] addition to the medium of the XylS effector 3MBz, in which case the activating agent is solely the complex XylS-3MBz, or [ii] supplementing the same medium with *m*-xylene, which is metabolically converted inside cells to 3MBz (the *bona fide* intracellular agonist of XylS) by the action of the *upper* TOL pathway enzymes (Figure 1). In this last scenario, *Pm* is activated by XylS-3MBz as well as by overproduction of the same TF (*m*-xyl triggers the *Ps* promoter for XylS expression, Figure 1). Although the Pm/XylS node of the TOL network is often abstracted as a binary switch with only ON/OFF states, the unique noisy nature of its output dashes this ideal vision and highlights the role of signal variability [26]. Figures 2A and 2B show flow cytometry results of promoter activity as measured in a variant of the natural TOL plasmid-bearing strain *P. putida* mt-2 called *P. putida* mt-2-Pm. This strain (Table 1) derives from the natural *P. putida* mt-2 isolate bearing the TOL plasmid pWW0, but it has been engineered to bear in its chromosome a transcriptional *Pm-GFP fusion. P. putida mt-*2*-Pm* thus it has [i] the sole source of XylS in the cognate pWWO-encoded gene, which is expressed through the *Ps* promoter borne by the TOL plasmid (see Figure 1 and Materials and Methods for details) and [ii] the encoded *Pm-GFP* in the chromosome. Because of this arrangement, the DNA region that encodes and supplies XylS transcriptional regulators is non-adjacent to the target promoter, from which it is physically separated in *trans.* The data of Figure 2 expose how different the expression regimes of *Pm* are depending on whether the XylS/Pm regulatory node is induced with *m*-xyl or 3MBz in this strain. Specifically, induction with *m*-xyl (and thus XylS overproduction and intracellular production of 3MBz) leads to an expression scenario where the noise range of the output signal allows a null overlap between the ON and the OFF states. In contrast, induction with externally added 3MBz (and low XylS) produces a remarkably different ON state while leaving the OFF state unaffected. Moreover, the noise regime brought about by added 3MBz consisted of a long flat-like distribution that went from the lowest intensity value to the highest. Output ON signals are thus patently different in either case, suggesting they originate in a different type of TF-promoter interplay beyond mere randomness. The ensuing question is whether we can interpret the noise patterns of Figure 2 on the background of the TOL network (Figure 1) and the two ways of activating *Pm* mentioned above. As shown below, this issue can be addressed by combining various modeling approaches with *ad hoc* experimentation.

**Figure 2:**
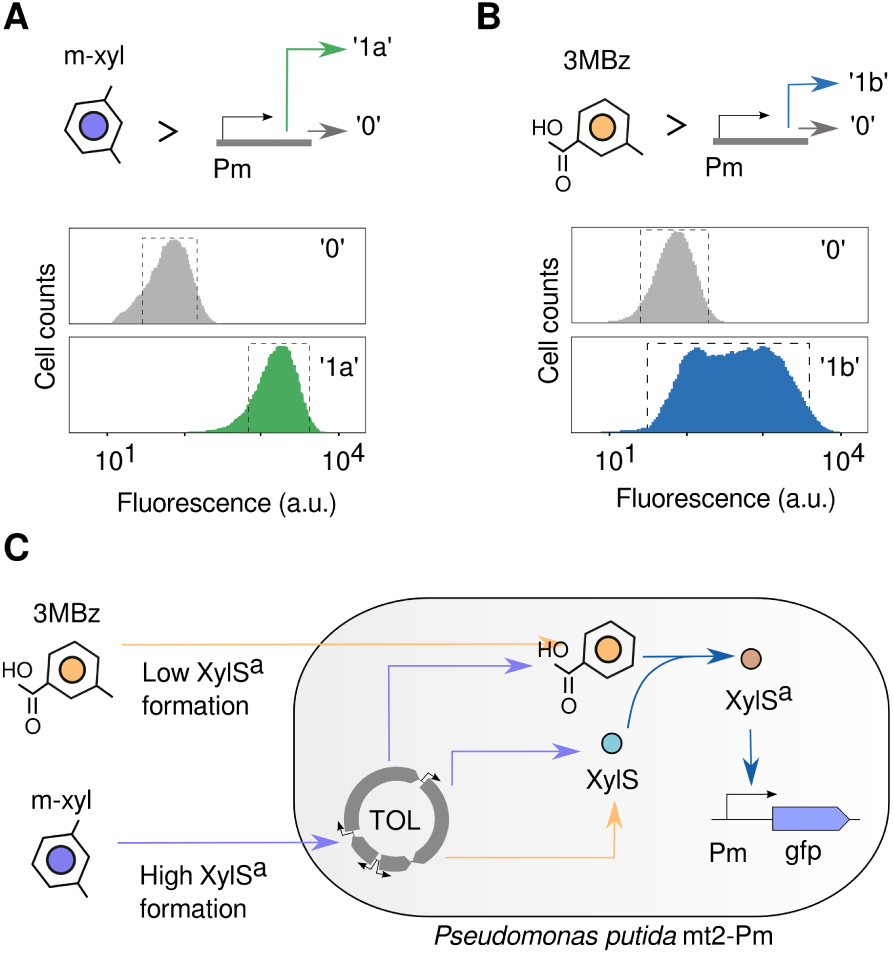
Variable noise patterns depending on input signal in P. putida mt-2-Pm strain. **A.** When the cells are subject to the presence or absence of *m*-xyl (*m*-xylene), the *Pm* promoter activity recorded (based on green fluorescent protein intensity) can be abstracted as binary switch with a 1 or ON state and a 0 or OFF state. Flow cytometry results show this behaviour, where the noise range allows a null overlap between the 0 and the 1 (called 1a to make a difference with the following). **B.** Using 3MBz (3-Methylbenzoate) as the inducer provokes again a switch-like behaviour in *Pm,* with 1/ON and 0/OFF states. In this occasion, as seen in the cytometry results, the noise range is much wider, going from the maximum expression to the minimum (ON state therefore called 1b). **C.** In this set of experiments, XylS molecules are produced by the TOL plasmid borne by the P. *putida* mt-2*-Pm* while the target Pm-GFP reporter fusion is inserted in the chromosome (see Materials and Methods), i.e. the source of the transcriptional factor and its target promoter are non-adjacent and encoded in separate monocopy (i.e. TOL plasmid and chromosome) replicons.

**Table 1:** Bacterial strains and plasmids used in this work

| Bacterial strain or plasmid | Relevant characteristics | Reference or source |
| --- | --- | --- |
| **Strains** |  |  |
| *Escherichia coli* |  |  |
| CC118λpir | Cloning host; $\Delta$(*ara-leu araD* Δ*lacX174 galE galK phoA thiE1 rpsE rpoB*(Rif) *agrE*(Am) *recA1* λpir lysogen | [51] |
| HB101 | Helper strain; F<sup>-</sup> λ<sup>-</sup> *hsdS20*(r<sub>B</sub><sup>-</sup> m<sub>B</sub><sup>-</sup>) *recA13 leuB6*(Am) *araC14* $\Delta$(*gpt-proA*)62 *lacY1 galK2*(Oc) *xyl-5 mtl-1 thiE1 rpsL20*(Sm<sup>r</sup>) *glnX44*(AS) | [52] |
| *Pseudomonas putida* |  |  |
| KT2440 | Wild-type strain; mt2 derivative cured of the TOL plasmid pWWO | [53, 54] |
| mt-2 | Wild-type strain bearing pWWO plasmid | [55] |
| mt-2-*Pm* | Gm<sup>r</sup>. *P. putida* KT2440 inserted in its genomic *attTn7* with the hybrid mini-Tn7 delivered by plasmid pBG-*Pm* (Supplementary Figure S6) | This work |
| KT-BGS | Gm<sup>r</sup>. *P. putida* KT2440 inserted in its genomic *attTn7* with the hybrid mini-Tn7 delivered by plasmid pBGS (Supplementary Figure S6) | This work |
| **Plasmids** |  |  |
| pRK600 | Cm<sup>r</sup>. Helper plasmid used for conjugation; *oriV* ColE1, RK2(*mob*<sup>+</sup> *tra*<sup>+</sup>) | [56, 57] |
| pTnS-1 | Ap<sup>r</sup>, *oriR6K*, *TnSABC+D* (Tn7 transposase) operon | [58] |
| pBG | Km<sup>r</sup>, Gm<sup>r</sup>, *oriR6K*, mini-Tn7 delivery vector; Tn7L and TnR bracketing a mobile DNA segment for engineering standardized BCD2-*msf GFP* reporter fusions (Supplementary Figure S6) | Zoebel *et. al.* (submitted) |
| pBG-*Pm* | Km<sup>r</sup>, Gm<sup>r</sup>, *oriR6K*. pBG inserted with *Pm* promoter and thus bearing a standardized *Pm-GFP* fusion | This work |
| pSEVA228 | Km<sup>r</sup>, *oriRK2*, *xylS/Pm* expression system | [59] |
| pBGS | Km<sup>r</sup>, Gm<sup>r</sup>, *oriR6K*, pBG inserted with the *xylS/Pm* module of pSEVA228 and thus bearing the TF adjacent to the same standardized *Pm-GFP* fusion as in pBG-*Pm* | This work |

### Noise deconvolution and rate optimization

The functioning of the regulatory node under inspection i.e. the regulator-promoter pair XylS-Pm, can be modelled following the kinetic rates shown in Figure 3A. The regulator in its active form, XylS^*a*^, binds *(k*_1_) *Pm* to fire its activity and unbinds *(*k*_–1_*) it back to the default, silent state. When bound, mRNA molecules are transcribed (*k*_2_) from the downstream gene, GFP, which produce proteins through translation (*k*_3_). Even when the regulator is not bound, there is some leak of basal *Pm* transcription (*k*_6_). To complete the model we include degradation rates for both mRNA (*k*_4_) and GFP (*k*_5_). Our goal here is to find those values for the rates that allow *Pm* to handle different noise regimes depending on the inducer used.

**Figure 3:**
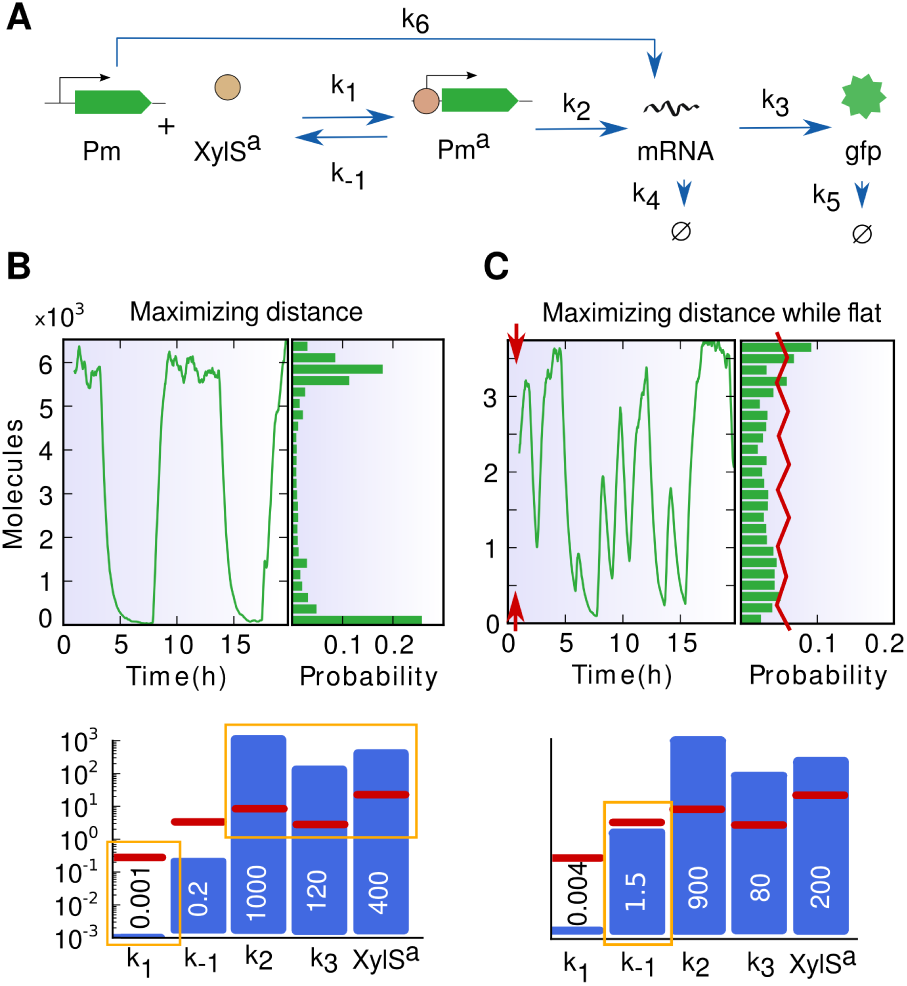
Rate optimization according to output-state fitness. **A.** Promoter (Pm) being studied in this work and the rates involved in the model. XylS^*a*^ is the activator of the inducible promoter in its active form. The complex *Pm*^*a*^ makes reference to the promoter with the regulator bound. Rates *k*_1_ and *k*_–1_ correspond to binding and unbinding events respectively. Transcription: *k*_2_. Translation: *k*_3_. Degradation rates: *k*_4_ and *k*_5_. Basal transcription rate is represented by *k*_6_. **B.** Rate optimization if terms of maximizing the distance between the maximum and minimum output level results in a bimodal distribution, where most cells are in either ON or OFF states and very few in-between at a given time. Bar-plot shows the optimization outcome for rate values. A very low binding rate guarantees a persistent OFF state while low unbinding value allows the high expression, as the regulator stay bound for longer periods. **C.** By forcing flatness (as well as amplitude) in the distribution, the noise profile has a smaller output range, marked by red arrows in the time-course graph. As seen in resulting rates, binding and unbinding values are increased to promote exploring intermediate expression values by boosting affinity instability. Most importantly, regulator numbers are drastically lowered, placing this value at the core of promoter activation. In both **A.** and **B.**, yellow markers in bar-plots highlight the most influential rates responsible for each behaviour. Red lines denote *standard values* (often used in the literature for computational analysis, see Materials and Methods.)

To this end, we consider a training vector *θ* with the rates that are mainly responsible for *Pm* dynamics defined as follows: θ = (*k*_1_, *k*_–1_, *k*_2_, *k*_3_, *[XylS^*a*^*]), where [x] denotes concentration of molecule x. Basal activity and degradations are specified within standard ranges found in the literature for mathematical analysis [27–31] (see Materials and Methods). In order to look for the set of values that best simulate the experimental output we describe two fitness parameters based on the ON state produced by 3MBz (Figure 2B): wide-range signal (*f*_1_) and flat-like surface (*f*_2_). When the first condition is applied to a series of simulations the optimized vector corresponds to *θf*_1_ = (0.001, 0.2,1000,120, 400), whose output is shown in Figure 3B. We can observe in the time-course plot that expression is either high or low with fast transitions in-between leading to a bimodal probability distribution. We then add the second fitness parameter to *θf*_1_: the probability distribution must have a flat-like surface. As this new condition has priority over the previous one, the optimization will output the vector that produces the widest range signal possible while assuring flatness. Figure 3C shows the simulation with the output vector *θf*_2_ = (0.004,1.5, 900, 80, 200) after the last optimization.

These results shed some light on the mechanics responsible for the ON state after induction with 3MBz. On the one hand, strong expression kinetics (transcription + translation) are needed to produce higher output values while a weak binding rate guarantees reaching the lower ones as the promoter remains empty for longer periods. On the other hand, it is the increment in the unbinding rate what helps generating the final flat-like distribution because it promotes affinity instability. Remarkably, the optimized number for the concentration of regulator molecules is rather low ([XylS^*a*^] = 200), matching our initial expectations since we are simulating a 3MBz induction case which should lead to fewer TF numbers than *m*-xyl.

In a further step, in order to check whether the noise regime observed in the ON state during *m*-xyl induction could be reproduced we increased the concentration of XylS^*a*^ molecules while leaving the rates of vector *θ*_*f*2_ untouched. Importantly, the simulations were successful at this stage and, as a result, this concentration is fixed at [XylS^*a*^] = 3000 molecules. These numbers produce the graph shown in Figure 4 (centre) in which the time-course lines at low and high induction correspond to the presence of 3MBz and *m*-xyl respectively. In supplementary Text S1 we detail the dynamics of the full TOL network and specify which kinetic values output the aforementioned XylS^*a*^ concentrations in each induction pathway. The balanced relationship of the two quantities, 200 and 3000, is based on previous qualitative observations [32].

**Figure 4:**
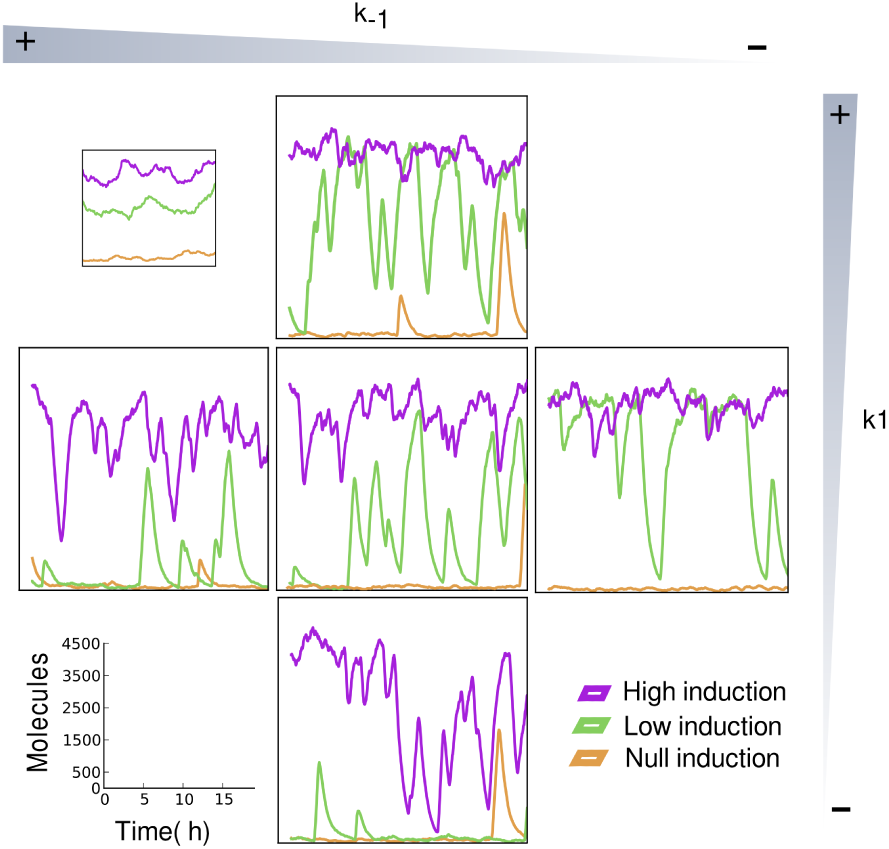
Effects of changes in binding/unbinding rates on *Pm* activity. Different time-course simulations are shown, where *Pm* promoter is exposed to three different concentrations of its regulator, XylS^*a*^: 10 (null induction, thus basal, yellow line), 200 (low induction, green line) and 3000 (high induction, purple line) molecules. The graph in the middle corresponds to the rates established in Figure 3C, with *k*_1_ =0.004 and *k*_–1_ = 1.5. Top: *k*_1_ reduced to 40% its original value. Bottom: *k*_1_ increased to 250%. Left: *k*_–1_ at 40%. Right: *k*_–1_ at 250%. Top-left graph shows the behaviour of a theoretical standard promoter with *k*_1_ =5 and k-_1_ =60, where the noise is proportional while the input increases.

During the course of this optimization process we observed that *Pm,* or its *in-in-silico* counterpart *θ*_*f*2_, is very specific. In other words: changes in certain rates can make the promoter to stop working correctly. Figure 4 tests the stability of the system under variable binding and unbinding rates. The graphs aligned vertically in Figure 4 share the value for *k*_–1_ while the value for *k*_1_ is increased to 250% its original value -upper graph- and decreased to 40% at the bottom. Equal ratios are applied to the changes in *k*_1_ within the horizontally lined up simulations. We clearly observe how the ideal behaviour is no longer maintained when those key rates are altered. We can then represent that scenario quantitatively by means of the differential variability (DV) of signals, a parameter that calculates changes in variance of expression. The middle scenario produces a DV= 11.7 while a standard promoter (up-left graph) has DV = 0.8 and any other scenarios show intermediate values between these two. Although these results allowed decoding the kinetic rates of *Pm*, they say nothing on the role played by the physical dynamics of the regulator-promoter interplay. To this end we adopted a separate approach as explained below.

### Dilution and extrinsic noise effects

Prior to decoding the information on regulator dynamics we tested the values of *θf*_2_ in a more computationally complex, although more realistic, scenario. As a result of simulating protein dilution dynamics and extrinsic noise, the previous optimized unbinding rate was decreased to maintain *Pm* functioning. For this we used our platform for simulating bacterial growth and population dynamics (DiSCUS, see Materials and Methods for details). In this agent-based simulation, each cell is a rod-shaped object that shows asynchronous growth and embodies its own copy of the mathematical model (reactions for Pm-GFP activity). Furthermore, this simulation incorporates creation and degradation rates for the regulator, which generates additional fluctuations. While intrinsic noise was modelled (as previously) by applying variability to the molecular species at stake, the streamlined simulation makes extrinsic stochasticity to affect the rates directly. In such spatial simulation, both kinds of noises are accumulated with time [33], but we introduced extrinsic fluctuations only when the cell divides into two newborn bacteria. This setup assumes a sort of idealized growth condition in which changes in the status of the cells are mostly due to division. On this basis, the fluctuations of a specific rate, r, are calculated according to its initial definition *r*_*0*_ by assigning a new value after each division that is randomly chosen within the range *r* ∈ [–*a* · *r*_0_, *a* · *r*_0_] where *a* = 0.2 (a 20% increase/decrease of the original rate value). Furthermore, molecular levels are lowered to half its value after division in order to simulate dilution [34]. With this setup we could then inspect the ON state produced by 3MBz induction, which is more sensitive to fluctuations in the number of regulators as these are present in fewer numbers (analysis output Shown in Supplementary Figure S1). In fact, Fig. S1 shows that due to the high variability in the concentration of XylS^*a*^, promoter output drops and the plateau behaviour is no longer maintained.

The question that arises on the basis of the above is what to change in *θ*_*f*2_ to restore normal functioning as observed in the experiments. The answer is given by the images of Figure 4: when the time-course line drops, a reduction of *k*_*–1*_ could help raising the levels of gene expression. We therefore changed unbinding rate to *k*_*–1*_ = 0.8. This single change gives equilibrium back to the system and the probability distribution moves drastically towards the sought flat-like shape. The explanation is that when we lower the unbinding rate we force the regulators to stay bound for longer to the promoter region. This then compensates the system for the strong fluctuations in XylS^*a*^, specially the inherent reduction in TF levels after division.

### System sensitivity to alterations in the concentration of the transcription factor

Simulations of the *Pm* response to gradual changes in regulator numbers reveals that previous estimated XylS^*a*^ figures for both ON states (200 and 3000 under 3MBz and *m*-xyl, respectively) are optimal to maximize differences between the two expression noise regimes. The simulated transfer function of *Pm*, shown in Figure 5A, indicates the range of the signal and its mean value at a given regulator concentration. Unlike other promoter transfer functions found in the literature [16,35–37] where transcriptional activity produces similar noise (error bars in graphs) regardless the input concentration, here the middle sector of the curve displays a very unique and wider noise profile. Taking a look in depth, it is indeed around 200 and 3000 XylS^*a*^ molecules where the noise ranges reach maximum and minimum levels respectively. The simulation of Figure 5B shows the system tested in continuous functioning where the inducer is changed sequentially. As observed, the output produced by *Pm* suggests a *trinary* (rather than *binary)* signal where there are three states, one OFF and two ON, each of them having a different shape that unequivocally recall their input. Population-based simulations (Supplementary Figure S2) with heterogeneous distribution of inducers over the surface where the cells grow on, emphasize the correlation between input compound and output signal in a visual fashion. It is also noteworthy this output corresponds to an amplitude-modulated (AM) signal that is produced from a frequency modulated (FM) promoter [38]. Therefore, there is a direct correlation between the time intervals of the bursting effect and the max-min distance (amplitude) of the resulting gene expression levels.

**Figure 5:**
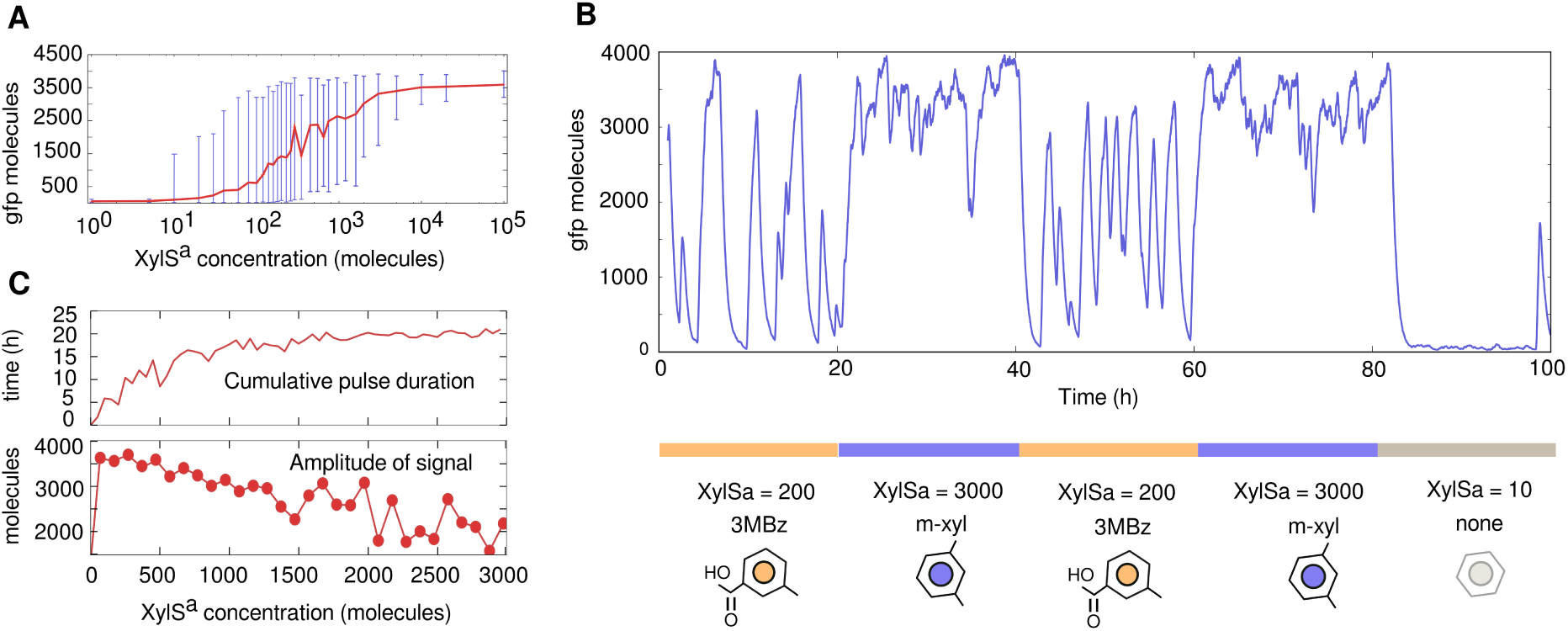
Analysis of *Pm* activity noise regarding regulator dynamics. Study of simulation sets where regulator concentration is the only parameter changed. A. Transfer function of *Pm*, which measures the output level resulting from different input concentrations. While the average value (red line) shows no relevant information but the usual, the noise produced by the signal (blue error bars, which denote max-min signal values) displays different behaviour according to input numbers, having wider range at middle concentrations. B. Concentration of GFP molecules over time while the input changes (3MBz, *m*-xyl or none). The three values of the logic displayed by the signal (called 0, 1a and 1b in Figure 2A-B) are: wide-range noise (times 0-20h and 40-60h), small-range high-level noise (times 20-40h, 60-80h) and small-range low-level noise (time 80h onwards). C 24h simulations with different XylS^*a*^ concentration each (from 0 to 3000 molecules) are use to measure: 1) the cumulative pulse duration, which corresponds to the length of time that Pm promoter is in the ON state (thus, its regulator is bound to the DNA) and 2) the amplitude of the signal, that is defined here as the distance (in molecules) from the highest value reached during the simulation to the minimum (measured in steady-state).

Two further analyses than link regulator dynamics with output noise are shown in Figure 5C. They must be interpreted on the background of the pulsing transcriptional bursts that frame activity of prokaryotic promoters as mentioned above. As shown in Figure 5C, two measurements, *cumulative pulse duration* and *signal amplitude,* were monitored in 24h simulations of the system at different XylS^*a*^ concentrations. The cumulative pulse makes reference to the core of the bursting effect, meaning the total time that *Pm* is in its active state, i.e. when XylS^*a*^ is bound to it. As a common trend, the total time of residence of XylS^*a*^ associated to *Pm* increases as the number of regulator molecules are higher. This points out the ON state produced during *m*-xyl induction to correspond to high numbers of XylS^*a*^. On the other hand, the amplitude of the signal (difference between the highest and the lowest values during each simulation) decreases except during the interval in which the number of XylS^*a*^ molecules is within the range ∈ [0..≃ 150], where the distance uppermost/lowermost signal increases. The cognate inflexion point thus can be explained as the number of XylS^*a*^ molecules that bring about the ON state during induction with 3MBz. This state of affairs is key to interpret noise in terms of physical TF-promoter dynamics, as explained below.

### Influence of intracellular regulator-promoter proximity on transcriptional output

On the basis of the above, we wondered whether the low-noise regime produced by *m*-xyl induction could be generated if 3MBz were the only input. When the model is interrogated with this question, the answer is positive when and only when 3MBz co-occurs with high numbers (3000) of the regulator XylS^*a*^. At a first glance this condition seems impossible to accomplish since the compound needed to stimulate production of more XylS^*a*^ molecules is *m*-xyl, not 3MBz (see Figure 1). But the situation changes if we rethink the possible effects of the number of XylS^*a*^ molecules produced in the physical matrix of a real bacterial cell. In the simulation scenario mentioned above it is assumed that each of the regulators is capable of binding the target promoter with a given, fixed rate as if it were a pure chemical reaction. In a real cellular setup (as the one adopted in Figure 2), one has to consider that due to imperfect diffusion caused by molecular crowding and non-homogenous micro-viscosity [39, 40] not all regulators are equally effective in reaching and binding cognate target DNA sequences. In reality, accessing the target promoter will be limited by the ease of diffusion towards the physical location where *Pm* is located in individual cells. To examine this possibility, we simulated individual protein trajectories [41–43] inside a cell, following a random Brownian motion [40,44,45].

Figures 6A-B record the trajectories of individual regulator molecules (XylS^*a*^) that are being expressed from what we call the *source* region. This location, that corresponds to the physical location of the *Ps* promoter from which *xylS* is expressed, can be abstracted as a defined, bounded site from which the XylS protein stems. Given the coupling of transcription and translation in the prokaryotic gene expression flow [46–48], it is safe to assume that the TF protein is produced in close proximity to the *Ps* promoter and the downstream *xylS* gene (Figure 1). But once produced, XylS^*a*^ must reach a physically separated *target* region where *Pm* is located in order to trigger transcription. This raises the different scenarios shown in Figures 6A-B, which diverge only in the number of proteins stemming from the source region. For a productive XylS^*a*^-promoter interaction to occur, the regulator must migrate from the place it is created towards the site where *Pm* is located. If the number of regulator molecules is low, there are necessarily *empty* locations inside the cell that XylS^*a*^ may not come across easily (at any one time). Should *Pm* be located in one of such regulator-empty sectors, it is physically impossible to materialize a productive contact, thus leading to an OFF promoter state. Scaling up this scenario to several thousand bacteria results in a situation such that each individual target region could accommodate a different number of regulators ranging from all to none, thus leading to a pronounced cell-to-cell variability. In contrast, when the regulatory proteins originated in the source region are abundant, there are hardly any empty areas inside the intracellular space and the variability range narrows. Note that the allocation of proteins at any given time does not necessarily match the distribution of trajectories (Figure 6B), meaning that the trails of the regulators are more space-dependent than their spreading. Due to the fact that regulators can bind and unbind several times their target promoter, it is their trajectories and not their final position that tell us which spatial regions are more likely to host TFs. As a result, the apparent binding rate of Figure 3, *k*_1_, in reality merges promoter-TF affinity proper (i.e. the physical interaction ability of the two molecular partners) with the occurrence of the regulator in the neighborhood of the promoter *in vivo*. Figure 6C shows the probability density of a simulated low-frequency regulator source placed at the middle of the cell compartment and the negative correlation between the distance d to this source and the density ρ of transcription factors. Therefore, the rate *k*_1_ of the original model can be replaced by a new function that is proportional to both parameters: intrinsic affinity and intracellular concurrence.

**Figure 6:**
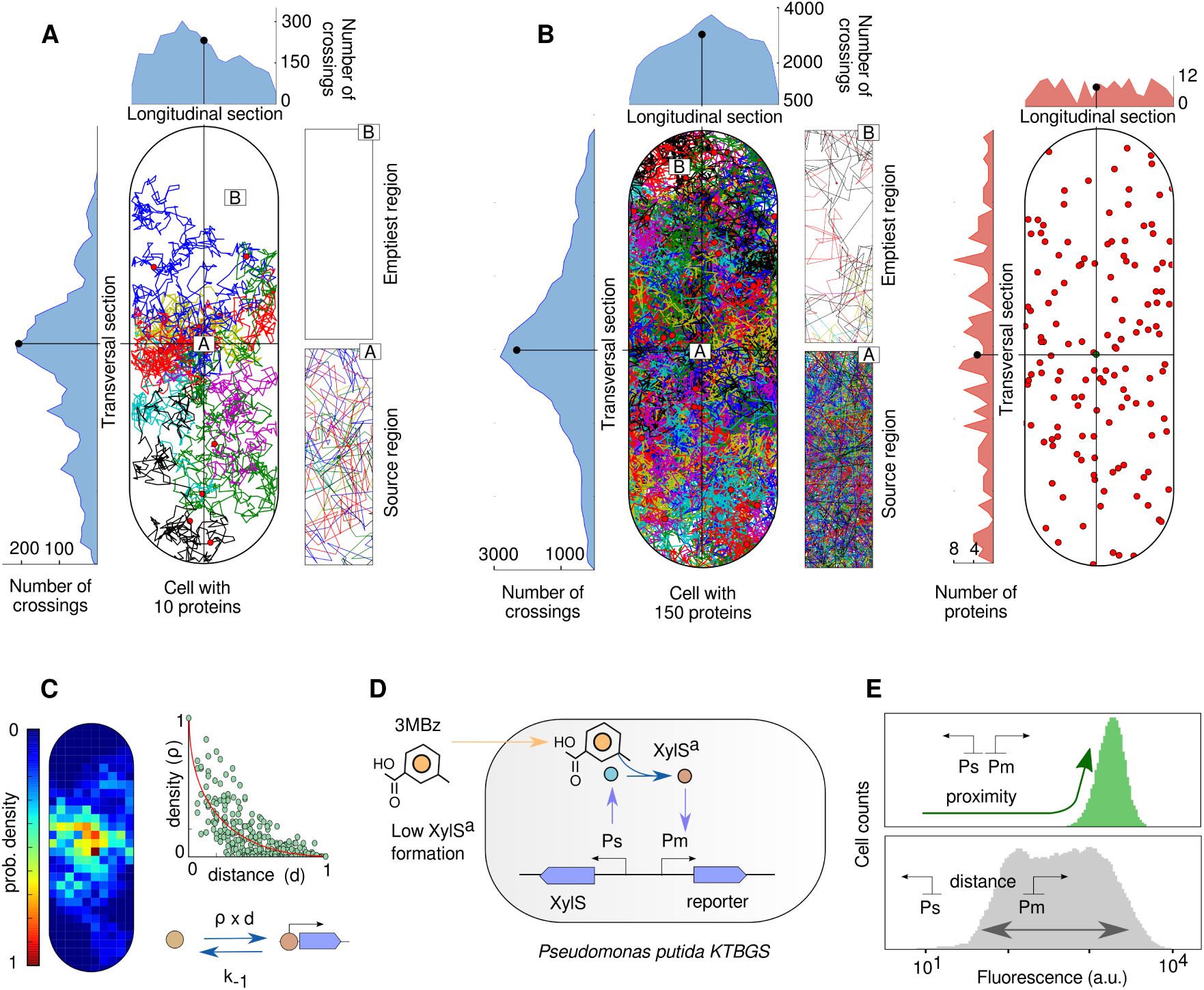
*Pm* promoter activity influenced by its distance from regulator source. **A** and **B.** Spatial distribution of simulated proteins following Brownian movement in a cell-like compartment. Each colour line inside the cell represents a trajectory of a protein from its source (placed at the middle of the space and labeled with the letter A) to its finally position at a given time. Density of trajectory positions at each section (longitudinal and transversal) is shown in sideways graphs; and two zoom-in regions (labeled A, B) with different trajectory points occupation are displayed in detail. **A.** Low frequency of protein production. **B.** High frequency of protein production. Picture on the right shows final protein position and highlights homogeneous distribution in sideways plots. **C.** Probability density per sub-compartment of the cells area according to a simulation with low frequency production reveals a strong negative correlation between distance from source and density of regulators. Binding rate *k*_1_, results in a combination of these two variables in a spatial scenario like the one considered. **D.** Physically re-arranged XylS/Pm regulatory node engineered in strain *P. putida* KT-BGS (Table 1) to maximize proximity between source (XylS production via *Ps* promoter) and target (Pm). Both promoters were inserted next to each other into the chromosome of strain KT2440, from where the TOL plasmid was removed (see Materials and Methods). **E.** Flow cytometry results with *P. putida* KT-BGS cells. As predicted by the model, using 3MBz as the inducer with minimal distance between source and target (upper graph, in green) gives the same results than using *m*-xyl with the reference *P. putida* mt-2-Pm strain (Figure 2A). Lower plot shows, as control, the 3MBz scenario in *P. putida* mt-2-Pm cells where source of the TF and the target promoter are not adjacent.

### Physical closeness between *Pm* and XylS decreases transcriptional noise

The experimental setting that reveals the differences in *Pm* noise caused by 3MBz -either added or generated metabolically from *m*-xyl- is one in which the promoter and the regulator are placed in somehow distant locations of the cell’s volume. This was brought about by placing the *Ps* promoter for *xylS* expression and the reporter Pm-GFP fusion in different replicons i.e., the TOL plasmid and the chromosome in *P. putida* mt-2-Pm, respectively (see Materials and Methods). One key projection of the model presented above is that physical proximity between the genomic sites bearing the *Ps-xylS* and Pm-GFP DNA segments should result in better occupation of *Pm* at lower XylS concentrations and therefore in a reduced GFP expression noise. To test this prediction we arrayed the *Ps-xylS* and Pm-GFP sequences within the frame of a mini-Tn7 transposon vector as explained in Materials and Methods. Delivery of such construct to the single *att*Tn*7* site of the strain *P. putida* KT2440 (i.e. same than *P. putida* mt-2 but free of the TOL plasmid) resulted in strain named *P. putida* KT-BGS (Table 1). In these engineered bacteria, the two components of the regulatory device under study (Ps→xylS/Pm→GFP) are thereby designed adjacent to each other, in monocopy and in a fixed chromosomal site. For the sake of the model, this means that distance between the TF source region and the promoter target region is artificially minimized (Figure 6D). In order to ensure that the modifications entered in P. putida KT-BGS do not distort the physical structure of the bacteria [49] we compared the apparent size and complexity of individual cells to those of its counterpart *P. putida* mt-2*-Pm* where *Ps-xylS* and Pm-GFP are separated. The data shown in Supplemental Figure S3 indicated that both strains are virtually indistinguishable as no significant differences were noticed in either physical quality. Once this was clarified, we repeated with strain *P. putida* KT-BGS the same induction experiment with 3MBz that was done previously with the reference *P. putida* mt-2-*Pm*. The flow cytometry results of this experiment are plotted in Figure 6E(up). For the sake of comparison, the lower plot of Figure 6E shows the same information of Figure 2 with strain *P. putida* mt-2- *Pm* added with 3MBz. Inspection of the data reveals that the proximity between *Ps-xylS* and Pm-GFP in *P. putida* KT-BGS results in a type of response to 3MBz that delivers a much narrower noise regime at high GFP intensity values. In this respect, the fluorescent signals of *P. putida* KT-BGS induced with 3MBz (where *xylS* expression is low but spatially proximal to *Pm*, Figure 6D) were indistinguishable to those of *P. putida* mt-2-Pm under *m*-xyl induction (TF expression high but distal to the target promoter). Moreover, the noise resulting from these two conditions diverge from the pattern observed in *P. putida* mt-2*-Pm* with 3MBz (low *xylS* expression from a source site separated from *Pm*). The mechanistic basis of the expression noise of each case is suggested by the model above (Figure 3), in particular our interpretation of rate *k*_1_. Under this frame (*k*_1_ = p*d), as the distance *Ps-xylS* to Pm-GFP decreases the density of the TF must be higher in order to keep the value of *k*_1_ constant as in the 1-dimensional simulations of Figure 5. In the case of strain *P. putida* KT-BGS, the proximity of *Ps-xylS* to Pm-GFP yields higher regulator numbers in the local molecular environment of *Pm*, what causes the same number of TFs to be available to its target promoter.

When the TFs were counted in the simulated target region we observed that a non-homogeneous intracellular space was needed in order to reach the optimized regulator numbers of Figure 5, highlighting the importance of different mobility areas in the cell [40]. Figure 7 shows the distribution of regulators within a cell-like compartment that is firstly *empty,* thus homogeneous (Figure 7A); and then compartmentalized by adding *low mobility* regions (Figure 7B) in which the Brownian motion is slowed down (see Materials and Methods). As observed, the difference in TF concentration caused by the proximity effect in the homogeneous space scenario (≃ 400 *vs.* 130 in Figure 7A) is not enough to reproduce the noise patterns *in-silico*. However, the addition of low mobility areas produces the accumulation of TFs within those regions and regulator numbers increase to reach the optimal proportion (≃ 3112 *vs.* 200 in Figure 7B). This latter scenario matches our experimental setup as the target region is inserted in the nucleoide, that corresponds to a highly condensed space. The noise-dependence of promoter-to-regulator distance is likely to be exacerbated if the TF at stake is very unstable, as seems to be the case with XylS [50]. Taken together, these data expose a new functionality to the intricate architecture of the regulatory network that governs biodegradation of *m*-xylene in *P. putida* mt-2.

**Figure 7:**
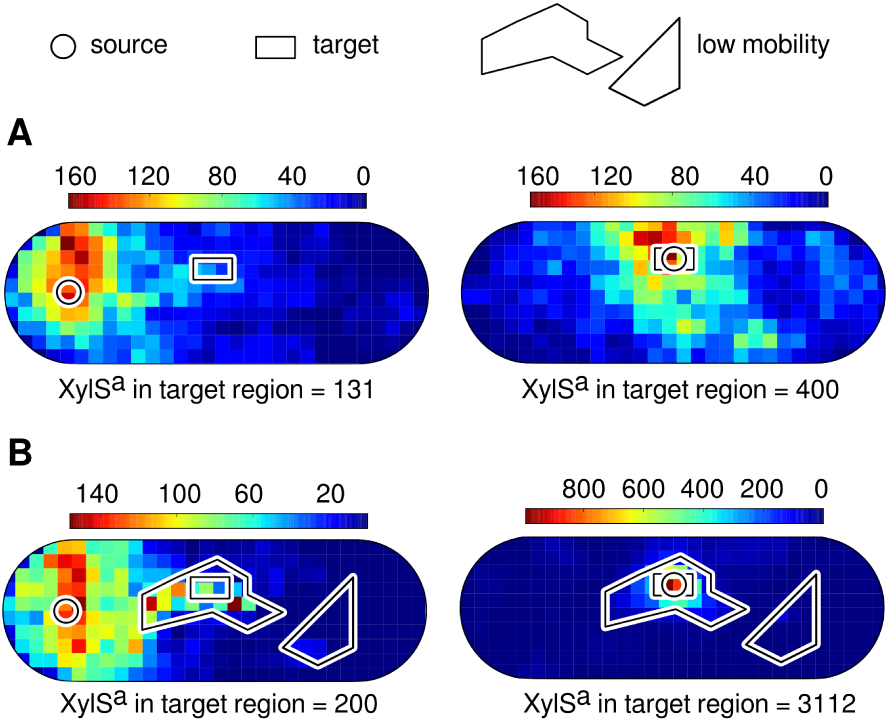
Local TF concentration affected by heterogeneous diffusion areas within the cell. Simulated regulators follow Brownian movement from a *source* region (circle) within a cell-like compartment. During the simulation, the number of regulators per iteration (scale bars) are monitored in squared sectors, with special attention to the *target* region (rectangle). A. In an homogeneous-space cell, the number of XylS^*a*^ molecules (trajectory points at a given time) within the target region corresponds to ≃ 130 when the source is not close (left) and ≃ 400 when both regions share the same local area (right). B. When running the simulation including *low mobility areas* (polygons), where TFs move slower, the number of XylS^*a*^ elements is ≃ 200 and ≃ 3000 in distant (left) and close (right) setups, respectively, matching the numbers optimized in the 1-dimensional analysis of Figure 5.

## Discussion

Intracellular signals are transmitted according to specific dynamics of the components involved in their transfer. Therefore, these communications are endowed with precise information, whose decodification promises valuable insights into cellular kinetic and structural properties [9]. It is signal variability, commonly referred to as gene expression noise [4,5], that constitutes the *fingerprint* of such a transmission, and thus the target data to interpret. In the case documented in this paper, the expression signals displayed initially by *Pm* promoter [19,22,23] activity in *P. putida* lead to highly-specific and stable noise patterns depending on what stimulus the cells were exposed to. Using mathematical modelling and computational analysis, we deconvoluted the flow cytometry data of each scenario to describe the kinetics that could reproduced that behaviour. As a result, the kinetic values that fits the experimental observations highlight the importance of the *bursting-specific* rates [13–16], binding and unbinding, where each of those values has its own influence in final promoter activity. Furthermore, and once the set of rates is established, we pinpointed how the dynamics of the *Pm*-regulator interplay determines gene expression output by entering spatial effects, in particular protein distribution within a cell. Our model, validated by the experiments shown above, accredits that the physical distance between the source of the regulator and the target promoter is translated into given noise patterns change drastically depending on promoter-TF proximity. This is due to the fact that regulators, or rather their trajectories (Figure 6) are not homogeneously distributed [60] and TFs are thus more likely to meet the promoters they regulate if located near the source [61]. This scenario was hypothesized by ten Wolde to explain the frequent genomic association of TFs and target promoters as an evolutionary remedy to an excess of noise [62,63]. In contrast, our analyses raise questions on whether gene expression noise caused by a non-homogeneous intracellular matrix s an adaptive trait in earnest which endows regulatory networks with emergent properties. In our case, we show that noise patterns of *Pm* can be altered by either changing the relative spatial positioning of the regulatory components (Figure 6) or their upstream kinetics (Supplementary Figure S4). As one of activity regimes of *Pm* is much more variable than the other, it may well happen that the noise-generating scenarios thereby described have been co-opted evolutionarily to create phenotypic heterogeneity within a population in order to increase its metabolic or else fitness [64,65]. This opens good opportunities to redesign heterologous expression systems for biotechnological purposes e.g. by decreasing phenotypic diversity of in a clonal population of producing cells [66]. Or just select the appropriate noise regime depending on the gene that is being expressed. For instance, if the gene of interest is a repressor (supplementary Figure S5) it is likely that we would want to reduce the noise. The data above also enter a new challenge in the engineering of non-native regulatory circuits or in, general, synthetic genetic implants in the genomic and biochemical chassis of a bacterial cell [36,67,68]. Every gene sequence and every protein (including TFs) may need a specific physical address in the 3D frame of a cell for an optimal performance, an issue that is hardly considered in contemporary Synthetic Biology and which surely deserves more attention. Finally, our results above put a new perspective on the much debated generality of transcription/translation coupling in prokaryotes [46–48], as the noise regime of given promoters is surely influenced by whether cognate TFs are generated close to the very genes which encode them or they have to migrate to other sites of the cell. These are all exciting questions that will be the subject of future studies.

## Materials and Methods

### Bacterial strains, growth conditions and genetic constructs

The bacterial strains and the plasmids used in this study are listed in Table 1. *Escherichia coli* cells were grown at 37° C in Luria Bertani (LB) medium and they were used as host for cloning procedures. *P. putida* cells were incubated at 30^°^C in M9 minimal medium supplemented with 2mM MgSO4 and 20 mM citrate as the sole carbon source [69]. When required, gentamycin (Gm 10 g mL^−1^), kanamycin (Km 50 g mL^−1^), ampicillin (Ap 150 g mL^−1^) and chloramphenicol (Cm 30 g mL^−1^) were added to growth media. Reporter strain *P. putida* mt-2*-Pm* is the original TOL plasmid pWW0-containing *P. putida* mt-2 which has been inserted in the single *att*Tn7 site of its genome with a *Pm*-GFP transcriptional fusion as follows. First, the *Pm* promoter sequence was amplified from plasmid pSEVA228 [59] as a 122 bp *Pad*/AvrII fragment with primers 5’TTAAT-TAAGGTTTGATAGGGATAAGTCC3’ and 5’CCTAG-GTCTGT TGCATAAAGCCTAA3’; and cloned into mini-Tn7 promoter-calibrating vector pBG (Zoebel *et al.,* submitted). The organization of this vector (Supplementary Figure S6A) is such that inserting promoter-bearing *Pac*I/*Avr*II originate an standardized translation/transcription fusion that minimizes any effect of the non-translated 5’ end of the reporter transcript in the final GFP readout. Cloning thereby the *Pm* promoter in pBG created mini-Tn7 delivery vector pBG-Pm (Supplementary Figure S6B). Second, this construct was mobilized to pWW0-containing *P. putida* mt-2 strain by tetra-parental mating as described in [57]. Finally, Gm^R^ exconjugants were verified for insertion of the hybrid mini-Tn7 transposon (carrying the Pm-GFP fusion) in an specific orientation at the *att*Tn7 site by amplifying the genomic region of interest with diagnostic PCR using primer pairs 5- Pput-glmS UP 5’AGTCAGAGTTACGGAATTGTAGG3’ / 3-Tn7L (5’ATTAGCTTACGACGCTACACCC3’ and 5- PpuglmS DOWN 5’TTACGTGGCCGTGCTAAAGGG3’ / 3- Tn7R 5’- CACAGCATAACTGGACTGATTTC3’. One of these clones yielding DNA products of 400 bp and 200 bp respectively [70,71] was designated as *P. putida* mt-2-Pm and used for the experiments discussed above. To obtain an entirely equivalent *P. putida* strain with a physically re-arranged XylS/Pm regulatory node, a 1088 bp DNA segment containing the array of regulatory parts — *xylS* ← *Ps* — *Pm ←* — was excised from plasmid pSEVA228 [59] as PacI/AvrII fragment and cloned in the corresponding sites of pBG vector (Supplementary Figure S6C). The resulting construct (pBGS) was mobilized to the genome of the pWW0-less strain *P. putida* KT2440 and Gm^R^ exconjugants tested for insertion of the mini-Tn7 transposon (with the Pm-GFP fusion adjacent to the *xylS* gene) in the same genomic site and orientation as before. One of these clones was named *P. putida* KT-BGS and picked for the tests of the effects of XylS/Pm closeness described in this paper. Note that this genetic strategy allowed a faithful comparison between the expression noise delivered by the Pm-GFP fusion borne by either *P. putida* mt-2*-Pm (Ps* → *xylS* and *Pm* in non-adjacent, separate replicons) and *P. putida* KT-BGS *(Ps* → *xylS* and Pm in close genomic proximity).

### Single cell analysis by flow cytometry

Single-cell experiments were performed with a Gallios (Perkin Elmer) flow cytometer. GFP was excited at 488 nm, and the fluorescence signal was recovered with a 525(40) BP filter. Strains grown overnight were diluted 1/100 and allowed to grow at 30° C in pre-filtered M9 citrate medium and incubated for 3-4 hours. After this pre-incubation, at the late-exponential phase (OD_600*nm*_=0.4), cells were treated with the inducer 3MBz at 1mM; cultures were then incubated with aeration at the temperature of 30°C, and at 3-hour induction, an aliquot of each sample was analyzed by flow cytometry. For every sample 20,000 events were analyzed.

### Biochemical reactions

The biochemical reactions that fully describe the model depicted in Figure 1 are the next:

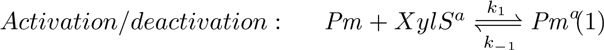

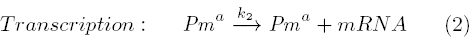

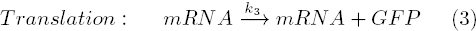

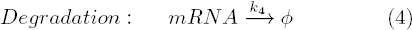

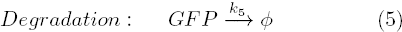

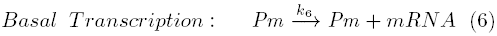

where *Pm*^*a*^ and *Pm* are the promoter with and without *XylS*^*a*^ bound -respectively-, *XylS*^*a*^ denotes the regulator in its active form, *mRNA* is the output of the transcription process and *GFP* is the final green fluorescent protein. The description of the rates is as follows: *k*_1_ is the binding rate of *XylS*^*a*^ to *Pm* (molecules^−1^ hour^−1^), *k*_–1_ the unbinding rate of *XylS*^*a*^ from the promoter (hour^−1^), *k*_*2*_ and *k*_*3*_ the transcription and translation rates (hour^−1^), *k*_*4*_ and *k*_*5*_ are the degradation rates of the *mRNA* and *GFP* (hour^−1^) and *k*_*6*_ the basal transcription of the promoter, which is the activity of *Pm* without regulator bound (hour^−1^). The full set of reactions and rates corresponding to the TOL network are shown in supplementary Text S1. The standard promoter referred to in the text corresponds to the following values: *θ* = (*k*_1_, *k*_*–1*_, *k*_2_, *k*_3_) ⇒ *θ* = (5, 60,150, 50). The values of Figure 3 where optimized within the ranges: *k*_1_ ∈ [0.001 – 1.2], *k_*_1_ ∈ [0.2 – 80], *k*_2_ ∈ [100 – 1000], *k*_3_ ∈ [10 – 120] and *XylS*^*a*^ ∈ [20 – 1500].

### Stochastic modelling of the biochemical network

Stochastic simulations are performed via Gillespie algorithm [72]. In this approach, unlike in a deterministic scenario, both chemical species and time are handled in a discrete fashion as well as the kinetic rates change their meaning to *probabilities* (of rates to be run). In order to represent the probability distribution corresponding to a time-course line of the expression level of a reporter, we consider each time point to be a single cell with its own molecule concentration. The Gillespie’s algorithm calculates that trajectory in asymmetric time lapses, *τ* = *–log(r)/a*_0_, were *r ∈* [0..1] is a random number and *a*_*0*_ the sum of reactions at that iteration. As a result the time intervals are variable in size and specially long when molecular levels are small, which means that during *τ* the system remains still. Thus, what the Gillespie’s algorithm returns needs to be converted into a time-course array where the time intervals are fixed, *τ* _α_, and are small enough to have cells (each time point) representing properly -in terms of frequency-all possible molecular levels. We use *τ*_*α*_ = 0.01h in this work (Figure S7). Differential variability (DV) is the relation between the variance of the noise under two different conditions. We measure it, as defined in [73], based on the expression 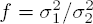 where 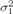 and 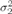 are the variances of the signal at low induction (3MBz) and high induction (*m*-xyl).

The inadequacy of deterministic approaches for the characterization of expression noise is shown in supplementary Figure S8.

### Population dynamics

We make use of our in-house software to simulate bacterial populations: DiSCUS (Discrete Simulation of Conjugation Using Springs - http://code.google.com/p/discus/) [74,75]. For the purpose of the present work, we include the management of extrinsic noise and protein dilution for an accurate simulation of the genetic noise. Furthermore, the next two extra rates are included in the model in order to force stochasticity in the formation of *XylS*^*a*^:

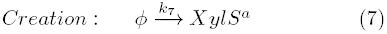

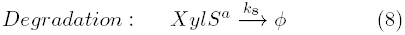

Extrinsic noise is simulated by changing the vector of rates after division. Thus, it reflects the fluctuations in the physical conditions of a *newborn* cell according to the initial set of rates. Being *θ*^*i*^ the vector of initial kinetic rates, every new vector affected by extrinsic noise (θ^*n*^) is described as,

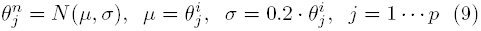

with *μ* being the mean, *σ* the standard deviation and *p* the dimension of the rate vector *θ*. For the standard deviation of the Gaussian noise we defined a 20% of the original value for every rate.

The simulated cells increase their dimension iteration-wise unless a threshold of pressure is reached (see DiSCUS website for details). That means that the stop condition determines the time during which the genetic circuit inside each cell will run for with the same rate vector. The doubling time is set initially to 450 iterations, where the rates inside each object run during 14 *simulated* seconds per iteration; so the *in-silico* gene is expressed during 105 minutes during the lifetime of the cell, according to the rates of Figure 3. The number of proteins is halved after division.

### Spatial protein movement

For the spatial intracellular simulation of regulators shown in Figures 6 and 7 we implemented a two-dimensional Brownian motion instance, written as an iteration scheme as follows:

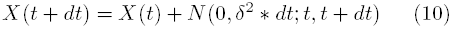

where *t* identifies the last time event, dt is the time step *(dt* = *T/N* with T the total time per iteration and N the number of steps, 15.0 and 1.0 respectively in our case) and δ the so called Wiener process parameter (0.25 for this study). Each protein runs during 400 iterations and the *slow mobility* regions change dt to 1.0 to simulate high density areas. All parameters are dimensionless and are set so that a simulated protein could cover the whole cell’s area (≂ 60 × 20 integer coordinates 2D lattice, where movement covers float numbers in between) during its lifetime. The scale-bars of Figure 7 measure the number of trajectory points (several for each protein) within the cell. Due to the fact that a regulator can bind and unbind different times its target promoter, it is trajectory points and not final protein numbers that are being monitored to match the 1-dimensional mathematical analysis. Therefore, there is a direct correlation between the number of TFs of Figure 5 and the trajectory points of Figure 7.

All computational simulation programs used (including the stochastic/deterministic algorithms/equations for network study, the population-based simulation software and the spatial motion functions) were written in Python (www.python.org) and run on a PC with debian (www.debian.org).

## Acknowledgements

This work was supported by the CAMBIOS Project of the Spanish Ministry of Economy and Competitiveness, the ST-FLOW, EVOPROG, ARISYS and EMPOWER-PUTIDA Contracts of the EU, and the PROMT Project of the CAM to VDL. Authors declare no conflict of interest.

## References

[1] Kaern M, Elston TC, Blake WJ, Collins JJ (2005) Stochasticity in gene expression: from theories to phenotypes. Nat Rev Genet 6: 451–64.

[2] Golding I, Paulsson J, Zawilski SM, Cox EC (2005) Real-time kinetics of gene activity in individual bacteria. Cell 123: 1025–1036.

[3] Raj A, van Oudenaarden A (2008) Nature, nurture, or chance: stochastic gene expression and its consequences. Cell 135: 216–226.

[4] Eldar A, Elowitz MB (2010) Functional roles for noise in genetic circuits. Nature 467: 167–173.

[5] Rinott R, Jaimovich A, Friedman N (2011) Exploring transcription regulation through cell-to-cell variability. Proceedings of the National Academy of Sciences 108: 6329–6334.

[6] McAdams HH, Arkin A (1999) Itsa noisy business! Genetic regulation at the nanomolar scale. Trends in genetics 15: 65–69.

[7] Hansen AS, O’Shea EK (2013) Promoter decoding of transcription factor dynamics involves a trade-off between noise and control of gene expression. Molecular systems biology 9.

[8] Munsky B, Neuert G, van Oudenaarden A (2012) Using gene expression noise to understand gene regulation. Science (New York, NY) 336: 183–187.

[9] Purvis JE, Lahav G (2013) Encoding and decoding cellular information through signaling dynamics. Cell 152: 945–956.

[10] Brehm-Stecher BF, Johnson EA (2004) Single-cell microbiology: tools, technologies, and applications. Microbiology and molecular biology reviews 68: 538–559.

[11] Czechowska K, Johnson DR, van der Meer JR (2008) Use of flow cytometric methods for single-cell analysis in environmental microbiology. Current opinion in microbiology 11: 205–212.

[12] Kortmann H, Blank LM, Schmid A (2011) Single cell analytics: An overview. In: High Resolution Microbial Single Cell Analytics, Springer. pp. 99–122.

[13] Taniguchi Y, Choi PJ, Li GW, Chen H, Babu M, et al. (2010) Quantifying E. coli proteome and transcriptome with single-molecule sensitivity in single cells. Science 329: 533–538.

[14] Zong C, So Lh, Sepúlveda LA, Skinner SO, Golding I (2010) Lysogen stability is determined by the frequency of activity bursts from the fate-determining gene. Molecular systems biology 6.

[15] Chong S, Chen C, Ge H, Xie XS (2014) Mechanism of transcriptional bursting in bacteria. Cell 158: 314–326.

[16] So Lh, Ghosh A, Zong C, Sepúlveda LA, Segev R, et al. (2011) General properties of transcriptional time series in *Escherichia coli*. Nat Genetics 43: 554–560.

[17] Nikel PI, Silva-Rocha R, Benedetti I, Lorenzo V (2014) The private life of environmental bacteria: pollutant biodegradation at the single cell level. Environmental microbiology 16: 628–642.

[18] Nikel PI, Martίnez-Garcίa E, de Lorenzo V (2014) Biotechnological domestication of pseudomonads using synthetic biology. Nature Reviews Microbiology 12: 368–379.

[19] de las Heras A, Fraile S, de Lorenzo V (2012) Increasing Signal Specificity of the TOL Network of *Pseudomonas putida* mt-2 by Rewiring the Connectivity of the Master Regulator XylR. PLoS Genet 8: e1002963.

[20] Nikel PI, Romero-Campero FJ, Zeidman JA, Goñi-Moreno Á, de Lorenzo V (2015) The Glycerol-Dependent Metabolic Persistence of *Pseudomonas putida* KT2440 Reflects the Regulatory Logic of the GlpR Repressor. mBio 6: e00340–15.

[21] Silva-Rocha R, Pérez-Pantoja D, de Lorenzo V (2013) Decoding the genetic networks of environmental bacteria: regulatory moonlighting of the TOL system of *Pseudomonas putida* mt-2. The ISME journal 7: 229.

[22] Ramos JL, Marqués S, Timmis KN (1997) Transcriptional control of the Pseudomonas TOL plasmid catabolic operons is achieved through an interplay of host factors and plasmid-encoded regulators. Annual review of microbiology 51: 341–373.

[23] González-Pérez MM, Marqués S, Dominguez-Cuevas P, Ramos JL (2002) XylS activator and RNA polymerase binding sites at the *Pm* promoter overlap. FEBS letters 519: 117–122.

[24] Pérez-Pantoja D, Kim J, Silva-Rocha R, Lorenzo V (2014) The differential response of the Pben promoter of *Pseudomonas putida* mt-2 to BenR and XylS prevents metabolic conflicts in m-xylene biodegradation. Environmental microbiology.

[25] González-Pérez MM, Ramos JL, Marqués S (2004) Cellular XylS levels are a function of transcription of xylS from two independent promoters and the differential efficiency of translation of the two mRNAs. Journal of bacteriology 186: 1898–1901.

[26] Goñi-Moreno A (2014) On genetic logic circuits: forcing digital electronics standards? Memetic Computing 6: 149–155.

[27] Dublanche Y, Michalodimitrakis K, Kümmerer N, Foglierini M, Serrano L (2006) Noise in transcription negative feedback loops: simulation and experimental analysis. Molecular systems biology 2.

[28] Goñi-Moreno A, Amos M (2012) Continuous computation in engineered gene circuits. Biosystems 109: 52 – 56.

[29] Balagaddé FK, Song H, Ozaki J, Collins CH, Barnet M, et al. (2008) A synthetic *Escherichia coli* predator-prey ecosystem. Mol Syst Biol 4: 187.

[30] Andersen JB, Sternberg C, Poulsen LK, rn SPB, Givskov M, et al. (1998) New Unstable Variants of Green Fluorescent Protein for Studies of Transient Gene Expression in Bacteria. Appl Environ Microbiol 6: 2240 – 2246.

[31] de Leon SBT, Davidson EH (2009) Modeling the dynamics of transcriptional gene regulatory networks for animal development. Developmental Biology 325: 317 – 328.

[32] Velazquez F, Parro V, de Lorenzo V (2005) Inferring the genetic network of *m*-xylene metabolism through expression profiling of the *xyl* genes of *Pseudomonas putida* mt-2. Molecular Microbiology 57: 1557–1569.

[33] Toni T, Tidor B (2013) Combined model of intrinsic and extrinsic variability for computational network design with application to synthetic biology. PLoS computational biology 9: e1002960+.

[34] Guido NJ, Wang X, Adalsteinsson D, McMillen D, Hasty J, et al. (2006) A bottom-up approach to gene regulation. Nature 439: 856–860.

[35] Moon TS, Lou C, Tamsir A, Stanton BC, Voigt CA (2012) Genetic programs constructed from layered logic gates in single cells. Nature 491: 249–253.

[36] Wang B, Kitney RI, Joly N, Buck M (2011) Engineering modular and orthogonal genetic logic gates for robust digital-like synthetic biology. Nature Communications 2: 508+.

[37] Bonnet J, Yin P, Ortiz ME, Subsoontorn P, Endy D (2013) Amplifying Genetic Logic Gates. Science 340: 599–603.

[38] Cai L, Dalal CK, Elowitz MB (2008) Frequency-modulated nuclear localization bursts coordinate gene regulation. Nature 455: 485–490.

[39] Miklos AC, Sarkar M, Wang Y, Pielak GJ (2011) Protein crowding tunes protein stability. Journal of the American Chemical Society 133: 7116–7120.

[40] Parry BR, Surovtsev IV, Cabeen MT, OHern CS, Dufresne ER, et al. (2014) The bacterial cytoplasm has glass-like properties and is fluidized by metabolic activity. Cell 156: 183–194.

[41] Elf J, Li GW, Xie XS (2007) Probing transcription factor dynamics at the single-molecule level in a living cell. Science 316: 1191–1194.

[42] Gahlmann A, Moerner W (2014) Exploring bacterial cell biology with single-molecule tracking and super-resolution imaging. Nature Reviews Microbiology 12: 9–22.

[43] Uphoff S, Kapanidis AN (2014) Studying the organization of DNA repair by single-cell and single-molecule imaging. DNA repair 20: 32–40.

[44] Uhlenbeck GE, Ornstein LS (1930) On the theory of the Brownian motion. Physical review 36: 823.

[45] Saffman P, Delbrück M (1975) Brownian motion in biological membranes. Proceedings of the National Academy of Sciences 72: 3111–3113.

[46] Gowrishankar J, Harinarayanan R (2004) Why is transcription coupled to translation in bacteria? Molecular microbiology 54: 598–603.

[47] Burmann BM, Schweimer K, Luo X, Wahl MC, Stitt BL, et al. (2010) A NusE: NusG complex links transcription and translation. Science 328: 501–504.

[48] Miller O, Hamkalo BA, Thomas C (1970) Visualization of bacterial genes in action. Science 169: 392–395.

[49] Weng X, Xiao J (2014) Spatial organization of transcription in bacterial cells. Trends in Genetics 30: 287–297.

[50] González-Pérez MM, Ramos JL, Gallegos MT, Marqués S (1999) Critical nucleotides in the upstream region of the XylS-dependent TOL meta-cleavage pathway operon promoter as deduced from analysis of mutants. Journal of Biological Chemistry 274: 2286–2290.

[51] Herrero M, de Lorenzo V, Timmis KN (1990) Transposon vectors containing non-antibiotic resistance selection markers for cloning and stable chromosomal insertion of foreign genes in gram-negative bacteria. Journal of bacteriology 172: 6557–6567.

[52] Boyer HW, Roulland-dussoix D (1969) A complementation analysis of the restriction and modification of DNA in *Escherichia coli*. Journal of molecular biology 41: 459–472.

[53] Nelson K, Weinel C, Paulsen I, Dodson R, Hilbert H, et al. (2002) Complete genome sequence and comparative analysis of the metabolically versatile *Pseudomonas putida* KT2440. Environmental microbiology 4: 799–808.

[54] Bagdasarian M, Lurz R, Rückert B, Franklin F, Bagdasarian M, et al. (1981) Specific-purpose plasmid cloning vectors II. Broad host range, high copy number, RSF 1010-derived vectors, and a host-vector system for gene cloning in Pseudomonas. Gene 16: 237–247.

[55] Worsey MJ, Williams PA (1975) Metabolism of toluene and xylenes by Pseudomonas (putida (arvilla) mt-2: evidence for a new function of the TOL plasmid. Journal of Bacteriology 124: 7–13.

[56] Kessler B, Herrero M, Timmis KN, De Lorenzo V (1994) Genetic evidence that the XylS regulator of the Pseudomonas TOL meta operon controls the Pm promoter through weak DNA-protein interactions. Journal of bacteriology 176: 3171–3176.

[57] Keen N, Tamaki S, Kobayashi D, Trollinger D (1988) Improved broad-host-range plasmids for DNA cloning in gram-negative bacteria. Gene 70: 191–197.

[58] Choi KH, Gaynor JB, White KG, Lopez C, Bosio CM, et al. (2005) A Tn7-based broad-range bacterial cloning and expression system. Nature methods 2: 443–448.

[59] Martίnez-Garcίa E, Aparicio T, Goñi-Moreno A, Fraile S, de Lorenzo V (2014) SEVA 2.0: an update of the Standard European Vector Architecture for de-/re-construction of bacterial functionalities. Nucleic acids research : gku1114.

[60] Ishihama A, Kori A, Koshio E, Yamada K, Maeda H, et al. (2014) Intracellular concentrations of transcription factors in *Escherichia coli* : 65 species with known regulatory functions. Journal of Bacteriolog : JB–01579.

[61] Dröge P, Müller-Hill B (2001) High local protein concentrations at promoters: strategies in prokaryotic and eukaryotic cells. Bioessays 23: 179–183.

[62] Warren P, Ten Wolde P (2004) Statistical analysis of the spatial distribution of operons in the transcriptional regulation network of *Escherichia coli*. Journal of molecular biology 342: 1379–1390.

[63] van Zon JS, Morelli MJ, Tnase-Nicola S, ten Wolde PR (2006) Diffusion of transcription factors can drastically enhance the noise in gene expression. Bio-physical journal 91: 4350–4367.

[64] de Jong IG, Haccou P, Kuipers OP (2011) Bet hedging or not? A guide to proper classification of microbial survival strategies. Bioessays 33: 215–223.

[65] Veening JW, Smits WK, Kuipers OP (2008) Bistability, epigenetics, and bet-hedging in bacteria. Annu Rev Microbiol 62: 193–210.

[66] Delvigne F, Goffin P (2014) Microbial heterogeneity affects bioprocess robustness: Dynamic single-cell analysis contributes to understanding of microbial populations. Biotechnology journal 9: 61–72.

[67] Lou C, Liu X, Ni M, Huang Y, Huang Q, et al. (2010) Synthesizing a novel genetic sequential logic circuit: a push-on push-off switch. Molecular systems biology 6.

[68] Daniel R, Rubens JR, Sarpeshkar R, Lu TK (2013) Synthetic analog computation in living cells. Nature 497: 619–623.

[69] Abril M, Michan C, Timmis K, Ramos J (1989) Regulator and enzyme specificities of the TOL plasmidencoded upper pathway for degradation of aromatic hydrocarbons and expansion of the substrate range of the pathway. Journal of Bacteriology 171: 6782–6790.

[70] Schweizer HP (2001) Vectors to express foreign genes and techniques to monitor gene expression in Pseudomonads. current opinion in biotechnology 12: 439–445.

[71] Bao Y, Lies DP, Fu H, Roberts GP (1991) An improved Tn7-based system for the single-copy in-sertion of cloned genes into chromosomes of gram-negative bacteria. Gene 109: 167–168.

[72] Gillespie DT (1977) Exact stochastic simulation of coupled chemical reactions. The Journal of Physical Chemistry 81: 2340–2361.

[73] Ho JW, Stefani M, dos Remedios CG, Charleston MA (2008) Differential variability analysis of gene expression and its application to human diseases. Bioinformatics (Oxford, England) 24.

[74] Gonñi-Moreno A, Amos M, de la Cruz F (2013) Multicellular Computing Using Conjugation for Wiring. PLoS ONE 8: e65986.

[75] Gonñi-Moreno A, Amos M (2012) A reconfigurable NAND/NOR genetic logic gate. BMC Systems Biology 6: 126.

